# Spinal Cord Injury Enhances Lung Inflammation and Exacerbates Immune Response After Acute Lung Injury

**DOI:** 10.1101/2024.08.21.608879

**Authors:** Bradford C. Berk, Amanda Pereira, Velia Sofia Vizcarra, Christoph Pröschel, Chia George Hsu

## Abstract

The severity of spinal cord injury (SCI) is closely tied to pulmonary function, especially in cases of higher SCI levels. Despite this connection, the underlying pathological mechanisms in the lungs post-SCI are not well understood. Previous research has established a connection between disrupted sympathetic regulation and splenocyte apoptosis in high thoracic SCI, leading to pulmonary dysfunction. The aim of this study was to investigate whether mice with low-level SCI exhibit increased susceptibility to acute lung injury by eliciting systemic inflammatory responses that operate independently of the sympathetic nervous system. Here, we employed T9 contusion SCI and exposed mice to aerosolized lipopolysaccharide (LPS) to simulate lung inflammation associated with acute respiratory distress syndrome (ARDS). Twenty-four hours post-LPS exposure, lung tissues and bronchoalveolar lavage (BAL) fluid were analyzed. LPS markedly induced proinflammatory gene expression (SAA3, IRG1, NLRP3, IL-1beta, MCP-1) and cytokine release (IL-1beta, IL-6, MCP-1) in SCI mice compared to controls, indicating an exaggerated inflammatory response. Infiltration of Ly6G/C positive neutrophils and macrophages was significantly higher in SCI mice lungs post-LPS exposure. Interestingly, spleen size and weight did not differ between control and SCI mice, suggesting that T9 SCI alone does not cause spleen atrophy. Notably, bone-marrow-derived macrophages (BMDMs) from SCI mice exhibited hyper-responsiveness to LPS. This study demonstrated an increase in lung inflammation and immune responses subsequent to low-level T9 SCI, underscoring the widespread influence of systemic inflammation post-SCI, especially pronounced in specific organs like the lungs.

**KEY POINTS:**

- Spinal cord injury leads to increased lung permeability following LPS inhalation.
- Spinal cord injury amplifies cytokine release after exposure to LPS.
- Spinal cord injury affects bone marrow-derived macrophages, showing heightened glycolytic activity and inflammatory conditions.

## Introduction

Individuals with spinal cord injury (SCI) face an elevated risk of respiratory complications, with acute lung injury (ALI) and acute respiratory distress syndrome (ARDS) emerging as prominent contributors to morbidity and mortality in this population [1, 2]. Notably, pneumonia is a primary contributor to premature mortality in SCI patients, even when considering factors such as immobility, aspiration, and ventilator utilization [3]. Importantly, it serves as an independent risk factor contributing to unfavorable neurological and functional long-term outcomes [3].

The underlying reasons for the increased susceptibility to respiratory challenges in SCI patients are multifaceted. First, high-level SCI results in changes in central nervous system (CNS)- immune system interactions[4, 5]. Specifically, disrupting the hypothalamic-pituitary-adrenal axis that regulates sympathetic nervous system (SNS) activity causes a hyperadrenergic state[6]. Increased catecholamines due to stress-related noradrenergic overactivation decrease blood flow to the abdominal organs, especially the spleen, causing splenic atrophy. Furthermore, high-level SCI stimulates the production and secretion from the adrenal gland of glucocorticoids (GCs) that promote apoptosis of immune cells, particularly lymphocytes. As a result, the incidence of infections is higher in the SCI population, particularly those with severe, high-level SCI, than in the able-bodied population[7–10]. Second, increased local as well as systemic inflammation serves as a catalyst for ALI and ARDS. The initial phase of SCI, occurring within a few hours, is characterized by the release of myelin debris, ATP, Ca^2+^, intracellular proteins, and mitochondrial DNA from damaged cells. These triggers induce inflammatory responses from local microglia, astrocytes, and circulating immune cells[11]. The subsequent phase involves cytokines and activated immune cells that prolong inflammation within SCI lesions, exacerbating tissue damage and causing systemic inflammation[12, 13].

Prior investigations in mice have demonstrated that a high thoracic (T3) injury disrupts sympathetic regulation of lymphoid organs, leading to splenocyte apoptosis [4, 14], and impairs lung immunity[14] during the acute phase of spinal cord injury (SCI). However, the effects of T9 (lower level) SCI on lung immunity, in particular during the sub-acute and chronic phases post-SCI and in the absence of a hyperactivated sympathetic nervous system are less well understood[15–17]. While respiratory complications, including pneumonia, are prevalent during acute care, particularly among ventilator-dependent patients, respiratory diseases manifest in 30% of SCI patients throughout their lifetime, irrespective of SCI lesion type[18], but a formal accounting of the incidence of these respiratory diseases in individuals with SCI is currently unavailable. This study addresses this gap by establishing a robust animal model to investigate the impact of acute lung injury (ALI) on lung inflammation post-T9 SCI. Understanding the intricate mechanisms involved in SCI-induced respiratory complications is critical for advancing therapeutic strategies and improving acute and chronic outcomes for individuals with spinal cord injuries[19].

## Materials and Methods

### Reagents

<u>Medium:</u> Dulbecco’s modified eagle medium (MT-10-013-CV), Fetal bovine serum (10437-028), Sodium pyruvate (11360070), and Streptomycin/penicillin (15140-122) were from Thermo Fisher Scientific (MA, USA). <u>Chemical:</u> ABC-HRP conjugate (PK-6100), Citrate Buffer pH 6.0, 10x (21545). Cell Lysis Buffer (10X) (9803s) was from Cell Signaling Technology (MA, USA). DAPI Fluoromount-G (0100-20) was from SouthernBiotech (AL, USA). Lipopolysaccharides from Escherichia coli (L3129), Heparin (H3149-25KU), Protease inhibitor cocktail (P8340), and EDTA solution (BP24831), Ketamine (K0001) was from SMH Pharmacy (NY, USA). Normal goat serum (S-1000) were from Vector Laboratories (CA, USA). RTL lysis buffer (79216) was from Qiagen (Hilden, Germany). Saline (2F7123) was from Cardinal Health (OH, Ireland). Xylazine (005469) was from Bimeda (IL, USA). <u>Assay kit:</u> Bicinchoninic acid (BCA) kit (BCA-1) was from Pierce ( IL, USA), iScript™ cDNA Synthesis Kit (1708891) and iQ™ SYBR® Green Supermix (1708882) were from Bio-Rad Laboratories (CA, USA). MCP-1 (RUO 432704), TNFα (RUO 430904), IL-1β (RUO 432604) and IL-6 (RUO 431304) ELISA kits were from BioLegend (CA, USA). <u>Antibody:</u> Alexa Fluor 488 Goat Anti-rabbit IgG(H+L) highly cross-adsorbed (A-11034) was from Molecular Probes (OR, USA). Ly6G&6C antibody (550291) from BD Biosciences (New Jersey, USA)

### Animals

Female C57BL/6N mice were purchased from (Jackson Laboratory, 000664) aged 10 weeks were used in the experiments. (Victor Chang Cardiac Research Institute). All mice were housed in a specific pathogen-free facility and kept in a temperature-controlled room set to a light and dark cycle of 12 hours each. The mice had ad libitum access to standard mouse chow and water. All of the experiments were approved by the University Committee on Animal Use For Research (UCAR) at the University of Rochester and followed National Institutes of Health guidelines for experimental procedures on mice.

### Mouse Spinal Cord Contusion Injury

Mice aged 12 weeks were anesthetized with a combination of intraperitoneal ketamine and xylazine. A laminectomy was performed at the thoracic 9 (T9) level, exposing the dura mater. The spinal cord underwent moderate bilateral contusion using a force-defined injury device (65 kDyn, Infinite Horizon Impactor 400, Precision Systems and Instrumentation). The injury site was closed in layers, encompassing muscle and skin. A heating pad maintained body temperature at 37 °C during surgery and the initial recovery period. Buprenorphine SR was administered subcutaneously at the time of surgery. Cages were equipped with food and DietGel® Recovery at the bottom, and lactated Ringer’s solution was given i.p. for the first 5 days to ensure hydration. Manual bladder expression occurred every 12 hours post-SCI until spontaneous voiding resumed.

### LPS inhalation in vivo

Age-matched female mice with SCI and control mice were subjected to LPS inhalation. The mice were randomly positioned within stainless steel compartments located in a polycarbonate rectangular chamber, connected to a nebulizer on one side and a vacuum system on the other. The LPS exposure dosage and duration adhered to established parameters from prior research[20]. A solution of LPS (10mg/10mL saline) was aerosolized into the chamber for 30 minutes, while control mice received aerosolized saline solution. Following complete aerosolization, mice were removed from the chamber and returned to the room. It is important to note that whole-body inhalation exposures did not induce pain or distress in the mice, as confirmed by continuous monitoring during exposure.

### Bone marrow progenitor cell isolation and bone marrow-derived macrophage (BMDM) differentiation

BMDMs preparation was performed as previously described [21]. L929 conditioned media which contains the macrophage growth factor M-CSF, was prepared by culturing L929 cells (ATCC) in complete DMEM (Thermo Fisher Scientific, MT10013CV) supplemented with 10% FBS, and 1% penicillin and streptomycin for 10 days at 37°C, 5% CO_2_. The L929 conditioned media, was collected, filtered (Vacuum Filter/Storage Bottle System, Corning, 431153), and stored at −80 °C until required. For isolation of BMDMs, tibias and femurs were removed from both male and female mice and flushed with media using a 26-gauge needle. Bone marrow was collected at 500 x g for 2 min at 4 °C, resuspended with complete DMEM medium and filtered through a 70-μm cell strainer (VWR international, 10199-657). Bone marrow progenitor cells were cultured in 100 mm dishes for 6-7 days in 70% complete DMEM medium and 30% L929-conditioned medium. Fresh medium (5 mL) was added on day 3. BMDMs were collected by scraping in cold PBS containing EDTA (1 mM). After centrifugation, BMDMs were seeded into 12-well plates at a density of 1.6 × 10^5^ cells/well in DMEM and incubated overnight before use.

### LPS stimulation in vitro

To stimulate BMDMs, cultures were rinsed twice with serum and antibiotic-free DMEM medium. The medium was replaced with fresh with serum and antibiotic-free DMEM medium. Cells were stimulated with or without LPS (100 ng/mL) for 3 hours.

### Collection of BAL fluid and lung tissue

Mice were anesthetized with a ketamine/xylazine intraperitoneal injection before euthanasia via cervical dislocation. The upper part of the trachea was cannulated and lavaged with 1 mL followed by 0.5 mL of PBS supplemented with 1 mM EDTA. Bronchoalveolar lavage (BAL) fluid underwent centrifugation at 500 x g for 5 min at 4°C. The resulting cell pellets from BAL fluid were lysed, and the protein content of the resulting cell lysates was measured using a BCA kit. Cytokines and chemokines were analyzed in BAL fluids. The superior lobe was rapidly frozen in liquid nitrogen for quantifying gene expression and homogenized in RTL lysis buffer.

### Collection of lung tissue for immunofluorescent staining

Mice were euthanized via intraperitoneal injection of ketamine (130 mg/kg) and xylazine (8.8mg/kg). The lungs were perfused free of blood by gentle infusion of 10 ml PBS containing 1 mM EDTA through the right ventricle. Lungs were inflated with 2mL, PBS-equilibrated 4% formaldehyde (VWR International, PI28908). Lung tissues were removed, drop fixed in 4% formaldehyde, embedded in paraffin, cut into 5 μm sections, and mounted onto slides. Sections were deparaffinized before use. Sections were washed 3 times in PBS followed by antigen retrieval for 20 minutes with steam using 1X Citrate buffer (Millipore, 21545), pH=6.0. Sections were blocked in 10% normal goat serum (Vector Laboratories, S-1000) in PBS for 1 hour at room temperature followed by overnight incubation at 4°C with 1:500 Ly6G&6C antibody (BD Biosciences, 550291) in 2% normal goat serum in PBS overnight. After three washes with PBS, fluorescence-conjugated secondary antibodies (Molecular Probes, A-11034 or A-11081) 1:1000 were incubated for 1 hour at room temperature and followed by three washes with PBS. Nuclei were stained with DAPI-fluoromount-G (Southern Biotech, 0100-20). Fluorescent images were captured using a confocal microscope (Olympus BX51, Software: SPOT Imaging software advanced). Ly6G&6C and DAPI positive cells were quantified by using NIH ImageJ software.

### Cytokine assay

BAL fluid samples were cleared by centrifugation at 16k×g for 5 min and stored at −20 °C. MCP-1, IL-6, TNFα and IL-1β levels were measured by ELISA kits according to the manufacturer’s instructions.

### RNA extraction and Real-time PCR

Complementary DNA was generated from 0.1 μg (cell) or 0.5 μg (tissue) RNA through reverse transcription. Amplification reactions included a target-specific portion of the reverse transcription product and 1 μM forward and reverse primers in SYBR Green qPCR master mix. Fluorescence was analyzed using a CFX connect real-time PCR system. Gene expression was normalized with β-actin and an independent reference sample using the delta delta CT method. Specific transcript amplification was confirmed by melting curve analysis at the end of each PCR experiment. The primer sequences are provided below. Mouse β-actin (Forward: TTCAACACCCCAGCCATGT, Reverse: GTAGATGGGCACAGTGTGGGT);; Mouse MCP-1 (Forward: AGGTCCCTGTCATGCTTCTG, Reverse: TCTGGACCCATTCCTTCTTG); Mouse IL-1β (Forward: GAGTGTGGATCCCAAGCAAT, Reverse: ACGGATTCCATGGTGAAGTC); Mouse IL-6 (Forward: GAGGATACCACTCCCAACAGACC, Reverse: AAGTGCATCATCGTTGTTCATACA); Mouse TNFα (Forward: TCTTCTCATTCCTGCTTGTGG, Reverse: GGTCTGGGCCATAGAACTGA); Mouse HIF-1a (Forward: CCCATTCCTCATCCGTCA AATA, Reverse: GGCTCATACCC ATCAACTCA); Mouse IRG-1 (Forward: GCAACATGATGCTCAAGTCTG, Reverse: TGCTCCTCCGAATGATACCA); Mouse GLUT1 (Forward: GGTTGTGCCATACTCATGACC, Reverse: CAGATAGGACATCCAGGGTAGC); Mouse HK1 (Forward: GTGGACGGGACGCTCTAC, Reverse: TTCACTGTTTGGTGCATGATT). Mouse NLRP3 (Forward: TTCCCAGACACTCATGTTGC, Reverse: AGAAGAGACCACGGCAGAAG); SAA3 (Forward: TTTCTCTTCCTGTTGTTCCCAGTC, Reverse: TCACAAGTATTTATTCAGCACATTGGGA)

### Statistics

Comparisons between SCI and control groups after LPS exposure were analyzed by two-way ANOVA followed by post hoc T tests using Bonferroni correction for multiple comparisons. In vitro experiments were repeated as three independent procedures, with duplicate or triplicate wells averaged prior to statistical analysis. All data were presented as mean ± SEM. GraphPad Prism 10.0 was used for statistical analysis. P values were indicated as follow: * < 0.05, ** < 0.01, *** < 0.001, **** < 0.0001.

## Results

The extent of pulmonary function impairment varies based on the severity of spinal cord injury (SCI), with more pronounced dysfunction observed at higher injury levels. Recent research has demonstrated that a high thoracic (T3) injury disrupts crucial sympathetic regulation of lymphoid organs, leading to splenocyte apoptosis. [4, 14]. In the case of T9 SCI, there is no hyperactivation of the sympathetic nervous system, but the injury triggers systemic inflammatory responses [19]. Limited information is available regarding the impact of low SCI on immune defense mechanisms in the lungs. In this investigation, we conducted T9 contusion SCI using the Infinite Horizon Impactor[22]. Ten days post-SCI, mice were exposed to aerosolized lipopolysaccharide (LPS), a bacterial endotoxin known to initiate a pro-inflammatory response in mouse lungs. This LPS model mimics a natural route of entry into the host respiratory tract and replicates aspects of the inflammatory cascades associated with lung inflammation and acute respiratory distress syndrome (ARDS) in humans[20]. Therefore, it serves as a valuable model for studying the effects of SCI that may amplify these cascades and intensify the disease process. SCI and control mice were subjected to a 30-minute exposure to LPS within stainless steel compartments placed inside a polycarbonate rectangular chamber, while control mice received aerosolized saline solution[23].

At 24 hours post-exposure to LPS, lung tissues and bronchoalveolar lavage (BAL) fluid were collected. Initially, we assessed lung injury and the expression of proinflammatory cytokine genes in lung tissues. Following exposure to saline, there was no difference in proinflammatory cytokine gene expression between control and SCI mice. LPS significantly increased the expression levels of SAA3, IRG1, NLRP3, IL-1β, MCP-1, and TNFα mRNA in lung tissues from all animals. However, these elevations were significantly greater in lung tissues from SCI mice (with the exception of TNFα, Fig. 1A-F). Consistently, SCI alone did not elicit cytokine release when compared to control mice. LPS exposure also led to a markedly greater increase in IL-1β, IL-6, and MCP-1 release in BAL fluid in SCI mice compared to control mice (Fig. 2A-D). These findings suggest that SCI alone did not provoke prolonged inflammation in the local lung tissue. Instead, SCI amplifies lung inflammation following exposure to LPS.

**FIGURE 1.**
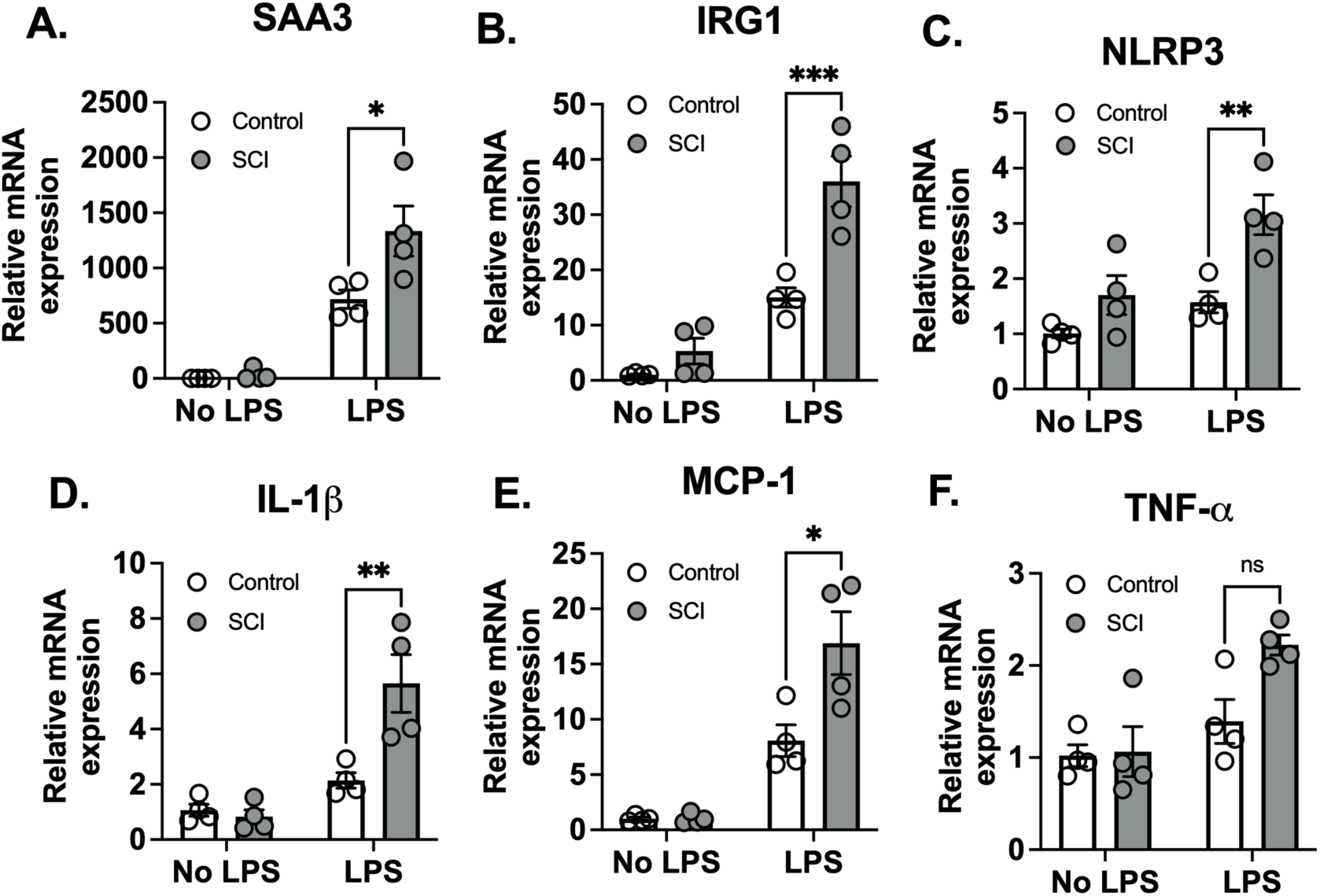
T9 SCI exaggerated lung inflammation after LPS exposure. T9 SCI exaggerated lung inflammation after LPS exposure. C57BL/6N mice underwent T9 SCI surgery. Ten days after SCI, mice were exposed to LPS (10mg/10mL saline) inhalation. Lung tissues were harvested at 24 hr after LPS exposure for gene expression of (A) SAA3, (B) IRG1, (C) NLRP3, (D) IL-1β, (E) MCP-1, and (F) TNF-α in the lung. (n=4 from each group). Data were expressed as mean ± SEM.

**FIGURE 2.**
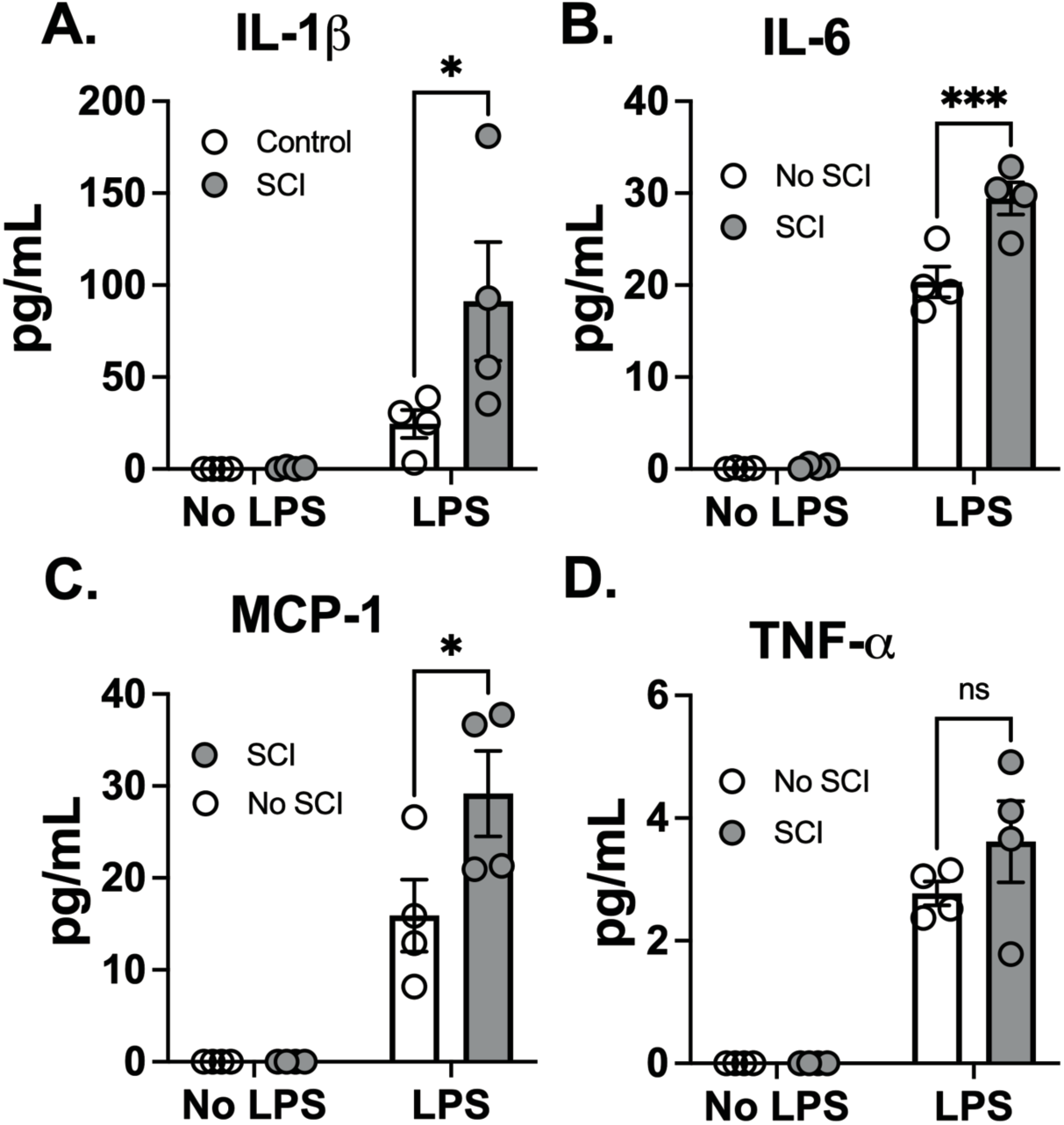
T9 SCI enhanced cytokine production after LPS exposure. C57BL/6N mice underwent T9 SCI surgery. Ten days after SCI, mice were exposed to LPS (10mg/10mL saline) inhalation. Bronchoalveolar lavage (BAL) fluids were harvested at 24 hr after LPS exposure for protein concentration in BAL fluid. Gene expression of (A) IL-1β, (B) IL-6, (C) MCP-1, and (D) TNF-α in the lung. (n=4 from each group). Data were expressed as mean ± SEM.

Inflammatory processes triggered by injury release various mediators such as cytokines, chemokines, and reactive oxygen species. These signaling molecules possess the capacity to influence the structural integrity of both the endothelial and epithelial barriers, thereby increasing lung permeability. Consequently, this phenomenon results in the effusion of fluids into the interstitial spaces and alveoli of the lung. Measuring lung permeability provides a quantitative measure of edema formation, which is a key pathological feature of acute lung injury[24, 25].

Ten days post-spinal cord injury (SCI), there was a two-fold increase in protein concentration in the BAL fluid, suggesting increased barrier permeability, although the difference was not statistically significant. Following exposure to LPS, protein levels in the BAL fluid markedly increased in SCI mice compared to control mice (Fig. 3A). Inflammatory cell activation and increased permeability form a positive feedback loop. Infiltrated cells release many mediators, that alter permeability, perpetuating inflammation and damaging endothelial barriers, correlating with the severity of lung injury. Inflammatory cell infiltration serves as a diagnostic marker for acute lung injury. Next, we investigated inflammatory cell infiltration using Ly6G/C staining, which marks neutrophils (Ly6G) and macrophages (Ly6C). No infiltration of Ly6G or Ly6C positive cells was observed in SCI mice following saline exposure. However, LPS inhalation led to a significant increase in Ly6G/C positive cells in lung tissue, indicating infiltration of neutrophils and macrophages. In accordance with cytokine findings, there was a significant elevation in Ly6G/C positive cells in lung tissue from SCI mice compared to control mice (Fig. 3B-C, arrows point to red cells). These results demonstrate that SCI enhances inflammatory cell infiltration and cytokine secretion in the lung after LPS exposure.

**FIGURE 3.**
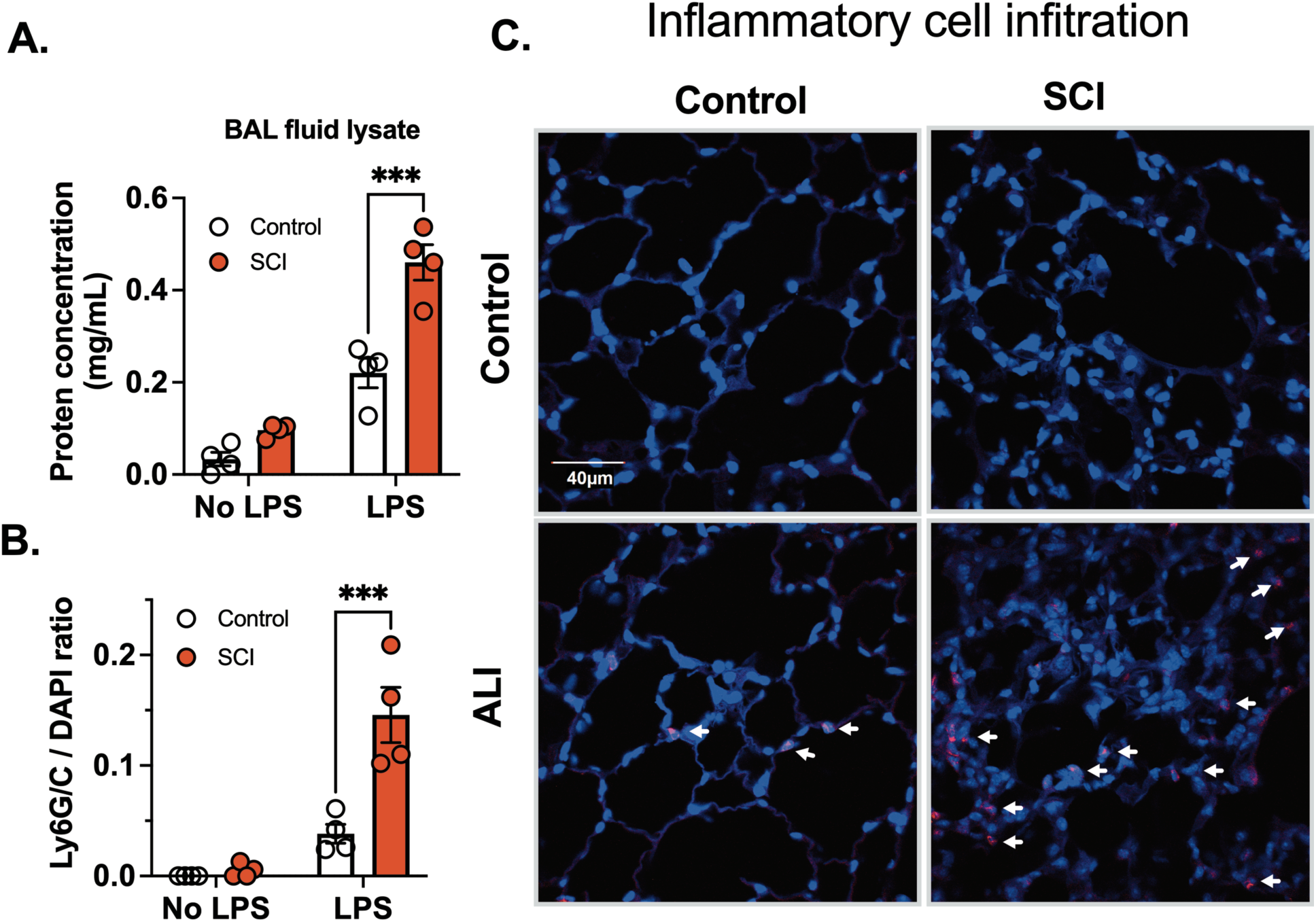
T9 SCI exaggerated inflammatory cell infiltration after LPS exposure. C57BL/6N mice underwent T9 SCI surgery. Ten days after SCI, mice were exposed to LPS (10mg/10mL saline) inhalation. BAL fluid and lung tissues were harvested at 24 hr after LPS exposure for (A) protein concentration in BAL fluid cell lysates. (B) Quantification results of Ly6G/C vs DAPI positive cells (C) Immunofluorescence of representative lung sections for Ly6G/C (Red) vs DAPI (Blue) (scale bar=40 µm). (n=4 from each group). Data were expressed as mean ± SEM.

Several studies have indicated that a high level SCI, as opposed to a low level SCI, leads to splenic atrophy and immune function suppression in mice, primarily due to exaggerated sympathetic reflexes[26–28]. To explore whether T9 SCI reduced spleen weights 10 days post-injury, we measured spleen size 24 hours after exposure to saline and LPS. However, we did not observe any significant differences in spleen weight or length between control and SCI mice (Fig. 4A-C). Furthermore, LPS exposure did not alter spleen weight or length compared to the saline groups (Fig. 4A-C). These findings suggest that T9 SCI alone is insufficient to induce splenic atrophy even in combination with acute lung injury.

**FIGURE 4.**
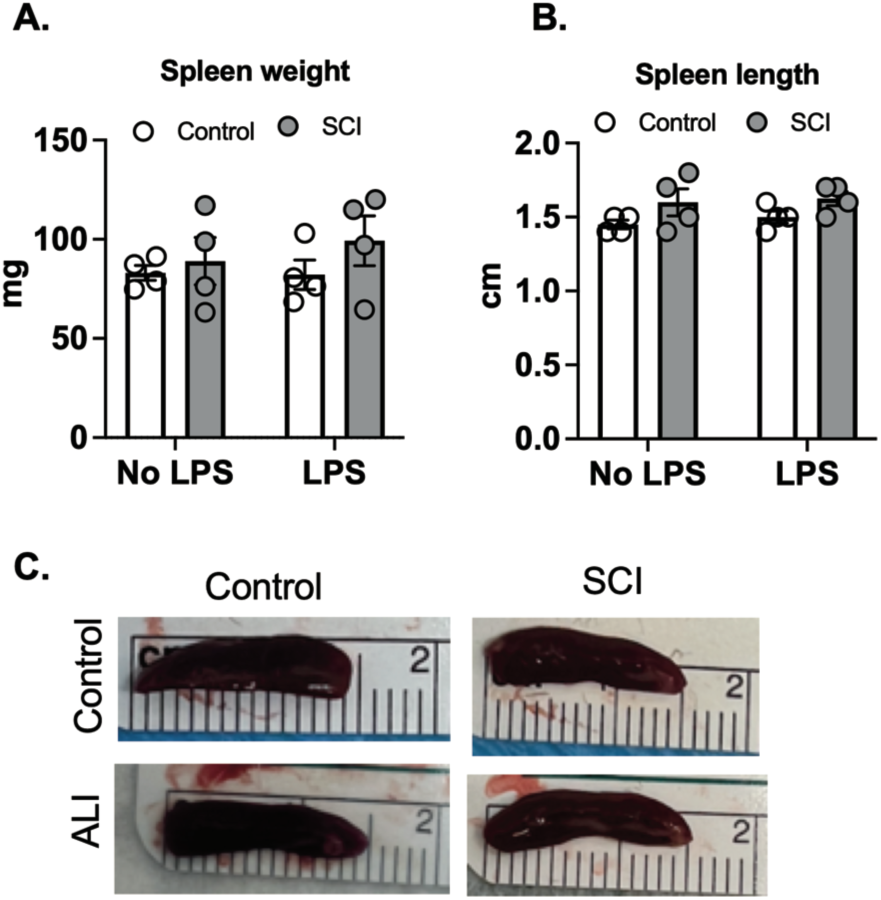
T9 SCI had no effect on spleen size after LPS exposure. C57BL/6N mice underwent T9 SCI surgery. Ten days after SCI, mice were exposed to LPS (10mg/10mL saline) inhalation. Spleens were harvested at 24 hr after LPS exposure. (A) Spleen weight (B) Spleen length was measured. (C) Representative spleen images for length (cm). (n=4 from each group). Data were expressed as mean ± SEM.

While previous studies emphasize mature leukocyte dysfunction as a hallmark of SCI-induced immune impairment, emerging evidence highlights the impact of SCI on the development and mobilization of immune cell precursors within the bone marrow. We collected bone marrow cells from mice 10 days post-SCI and from control mice, subsequently differentiating them into macrophages. In assessing the influence of SCI on these bone marrow-derived macrophages (BMDMs), we examined the expression of both glycolytic and inflammatory marker genes. Our findings indicate that BMDMs from SCI mice exhibit increased glycolytic activity and heightened inflammatory conditions (Fig. 5A). After subjecting these macrophages to LPS for 3 hours, BMDMs from SCI mice demonstrated a significantly increased response to LPS, as measured by mRNA levels of NLRP3 and IRG1 (Fig. 5B-C). These data show that SCI may alter the maturation of bone marrow derived macrophage precursors thereby contributing to a state of hyper-responsiveness in the lung.

**FIGURE 5.**
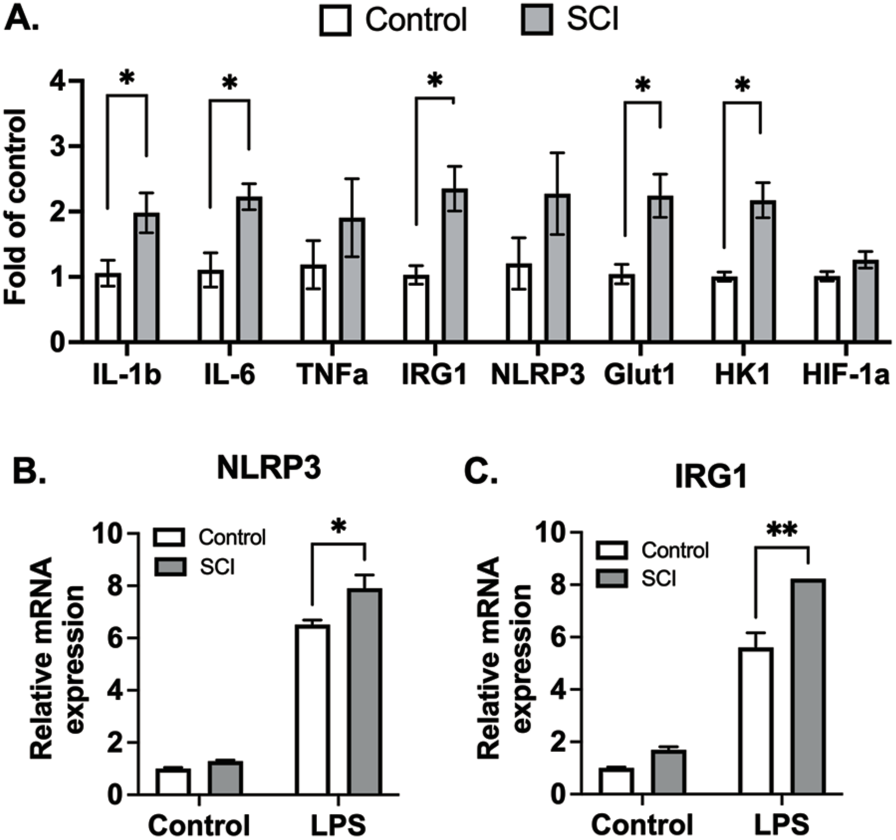
BMDMs from SCI mice are more glycolytic and responsive to inflammatory stimuli. C57BL/6N mice underwent T9 SCI surgery. Ten days after SCI, BMDMs were harvested and differentiated from control and SCI mice. (A) Basal gene expression of BMDMs. (B) NLRP3 and (C) IRG1 gene expression after LPS stimulation for 3 hr (n=4 experiments). Data were expressed as mean ± SEM.

## Discussion

In this study, our aim was to elucidate the impact of SCI, specifically at the T9 level, on pulmonary function and immune responses, particularly in the context of lung inflammation induced by aerosolized LPS. Our results indicate that ten days post-SCI recovery, mice exposed to LPS exhibited a marked increase in proinflammatory cytokine gene expression in the lung tissues compared to control mice. We observed a significant increase in cytokine release in BAL fluid from SCI mice compared to control mice following LPS exposure. Additionally, there was a significant increase, both at baseline and in response to LPS, in bone marrow-derived macrophages harvested from SCI mice, indicating that the acute increase in systemic inflammation is maintained in these macrophages even after the systemic inflammation disappears. These findings collectively indicate that SCI amplifies lung inflammation in response to LPS, emphasizing the role of SCI in exacerbating inflammatory cascades associated with lung inflammation, pneumonia, and acute respiratory distress syndrome.

The LPS exposure mouse model has been used to study mechanisms of human lung diseases and for the discovery of new drug targets for ALI[29, 30]. The LPS inhalation model provides a more physiological route of entry into the host respiratory tract compared to intratracheal or intranasal delivery. The utilization of our SCI-ALI mouse model offers a unique preclinical model for examining the impact of SCI on immune defense mechanisms and exploring potential avenues for drug development in the context of pulmonary dysfunction.

Furthermore, a significant elevation in protein concentration in BAL fluid from SCI mice following LPS exposure compared to control mice, suggests increased lung permeability. A positive feedback loop involving inflammatory cell activation and increased permeability was evident, with infiltrated cells releasing mediators that perpetuated inflammation and damaged barriers. This phenomenon correlated with the severity of lung injury, as demonstrated by increased inflammatory cell infiltration in SCI mice, marked by Ly6G/C staining. It is important to mention that within the first ten days after spinal cord injury (SCI), a slight rise in protein concentration in the BAL fluid was observed. This highlights the significant finding that there are acute changes in vascular damage and alterations in the lung microenvironment following acute SCI inflammation. Clinically, this implies that patients with SCI will have exaggerated immune responses to ventilator-associated pneumonias.

Interestingly, despite previous studies indicating splenic atrophy and immune function suppression in high-level SCI, our investigation into T9 SCI did not reveal significant differences in spleen weight or length between control and SCI mice, either with or without LPS exposure. Previous studies show that LPS treatment triggers splenomegaly and the accumulation of immune cells in the spleen four days later [31]. In our experiment, spleens were harvested 24 hours after acute lung injury (ALI). We selected this time point based on our prior discovery that lung permeability, inflammation, and immune cell infiltration reach their peak 24 hours after LPS exposure[20]. Despite this, our findings indicate that T9 SCI alone is inadequate to prompt alterations in spleen size even during the acute phase of acute lung injury, ten days post-injury.

Moving beyond the lungs, our study examined the impact of T9 SCI on immune cell precursors within the bone marrow. Analysis of bone marrow-derived macrophages (BMDMs) revealed heightened glycolytic activity and increased inflammatory conditions in BMDMs obtained from SCI mice. Similar outcomes were observed in the context of high-level C3 SCI, indicating hyperinflammatory responses in bone marrow cells by day 3 post-injury[32]. In humans, SCI has been shown to stimulate apoptosis and decrease proliferation of bone marrow cells[33, 34].

These cumulative findings suggest that SCI may directly affect immune function at the origin of all immune cells, specifically within the bone marrow cells. Expanding the focus beyond nerve injury, prior research has established associations between conditions such as diabetes, a western diet, as well as viral and bacterial infections in mice with elevated baseline inflammation in bone marrow cells [35–37]. This implies that sterile or non-sterile inflammation may exert a direct influence on regulating the bone marrow microenvironment. Subsequent research endeavors are crucial to unravel the mechanisms contributing to SCI-induced bone marrow hyperinflammation and determine whether such mechanisms are responsible for acute or chronic immune dysfunction.

In conclusion, T9 SCI causes a significant increase in inflammation stimulated by LPS inhalation 10 days after SCI when there is little systemic inflammation. Our data indicates that this heightened inflammatory memory is likely attributable to the impact on bone marrow cells, particularly those undergoing differentiation into macrophages. Various SCI injury levels distinctly affect pulmonary function, immune responses, and the bone marrow microenvironment. The observed rise in lung inflammation and enhanced reactivity of immune cell precursors post-SCI have significant implications for understanding systemic effects on immune function. These findings pave the way for future investigations and the development of SCI subtype-specific therapeutic interventions in this context.

## Declarations

### Ethics approval and consent to participate

Not applicable

### Consent for publication

Not applicable

### Availability of data and materials

All the data and materials are provided in the manuscript.

### Authors’ contributions

C.G.H. designed research; A.P., V.S.V., C.P., C.G.H. performed research; B.C.B., C.G.H. contributed reagents/analytic tools; V.S.V., C.G.H. analyzed data; C.G.H., B.C.B. wrote the paper; C.G.H., B.C.B. provided funding.

## Acknowledgements

None

